# Crowdsourced MRI quality metrics and expert quality annotations for training of humans and machines

**DOI:** 10.1101/420984

**Authors:** Oscar Esteban, Ross W Blair, Dylan M Nielson, Jan C Varada, Sean Marrett, Adam G Thomas, Russell A Poldrack, Krzysztof J Gorgolewski

**Affiliations:** Dept. of Psychology, Stanford University, CA, USA.; Data Science and Sharing Team, National Institute of Mental Health, Bethesda, MD, USA.; Functional MRI Facility, National Institute of Mental Health, Bethesda, MD, USA.

## Abstract

The neuroimaging community is steering towards increasingly large sample sizes, which are highly heterogeneous because they can only be acquired by multi-site consortia. The visual assessment of every imaging scan is a necessary quality control step, yet arduous and time-consuming. A sizeable body of evidence shows that images of low quality are a source of variability that may be comparable to the effect size under study. We present the MRIQC Web-API, an open crowdsourced database that collects image quality metrics extracted from MR images and corresponding manual assessments by experts. The database is rapidly growing, and currently contains over 100,000 records of image quality metrics of functional and anatomical MRIs of the human brain, and over 200 expert ratings. The resource is designed for researchers to share image quality metrics and annotations that can readily be reused in training human experts and machine learning algorithms. The ultimate goal of the database is to allow the development of fully automated quality control tools that outperform expert ratings in identifying subpar images.

## Background & Summary

Ensuring the quality of neuroimaging data is a crucial initial step for any image analysis workflow because low-quality images may obscure the effects of scientific interest^1–4^. Most approaches use manual quality control (QC), which entails screening every single image of a dataset individually. However, manual QC suffers from at least two problems: unreliability and time-consuming nature for large datasets. Unreliability creates great difficulty in defining objective exclusion criteria in studies and stems from intrinsically large intra-rater and inter-rater variabilities^5^. Intra-rater variability derives from aspects such as training, subjectivity, varying annotation settings and protocols, fatigue or bookkeeping errors. The difficulty in calibrating between experts lies at the heart of inter-rater variability. In addition to the need for objective exclusion criteria, the current neuroimaging data deluge makes the manual QC of every magnetic resonance imaging (MRI) scan impractical. For these reasons, there has been great interest in automated QC^5–8^, which is progressively gaining attention^9–16^ with the convergence of machine learning solutions. Early approaches^5–8^ to objectively estimate image quality have employed “image quality metrics” (IQMs) that quantify variably interpretable aspects of image quality^8–13^ (e.g., summary statistics of image intensities, signal-to-noise ratio, coefficient of joint variation, Euler angle, etc.). The approach has been shown sufficiently reliable in single-site samples^8,11–13^, but it does not generalize well to new images acquired at sites unseen by the decision algorithm^9^. Decision algorithms do not generalize to new datasets because the large between-site variability as compared to the within-site variability of features poses a challenging harmonization problem^17,18^, similar to “batch-effects” in genomic analyses^19^. Additional pitfalls limiting fully automated QC of MRI relate to the small size of databases that include quality annotations, and the unreliability of such annotations (or “*labels noise*”). As described previously, rating the quality of every image in large databases is an arduous, unreliable, and costly task. The convergence of limited size of samples annotated for quality and the labels noise preclude the definition of normative, standard values for the IQMs that work well for any dataset, and also, the generalization of machine learning solutions. Keshavan et al.^16^ have recently proposed a creative solution to the problem of visually assessing large datasets. They were able to annotate over 80,000 bidimensional slices extracted from 722 brain 3D images using BraindR, a smartphone application for crowdsourcing. They also proposed a novel approach to the QC problem by training a convolutional neural network on BraindR ratings, with excellent results (area under the curve, 0.99). Their QC tool performed as well as MRIQC^9^ (which uses IQMs and a random forests classifier to decide which images should be excluded) on their single-site dataset. By collecting several ratings per screened entity, they were able to effectively minimize the labels noise problem with the averaging of expert ratings. As limitations to their work, we would count the use of 2D images for annotation and the use of a single-site database. In sum, automating QC requires large datasets collected across sites, and rated by many individuals in order to ensure generalizability.

Therefore, the MRIQC Web-API (web-application program interface) provides a unique platform to address the issues raised above. The database collects two types of records: i) IQMs alongside corresponding metadata extracted by MRIQC (or any other compatible client) from T1w (T1-weighted), T2w (T2-weighted) and BOLD (blood-oxygen-level-dependent) MRI images; and ii) manual quality ratings from users of the MRIQC software. It is important to note that the original image data are not transferred to the MRIQC Web-API.

Within fourteen months we have collected over 50,000 and 60,000 records of anatomical and functional IQMs, respectively (Figure 1). These IQMs are extracted and automatically submitted (unless the user opts out) with MRIQC (Figure 2). Second, we leverage the efficiency of MRIQC’s reports in assessing individual 3D images with a simple interface that allows experts to submit their ratings with a few clicks (Figure 3). This assessment protocol avoids clerical errors from the operator, as ratings are automatically handled and registered. In other words, MRIQC users are building a very large database with minimal effort every day. As only the IQMs and manual ratings are crowdsourced (i.e. images are not shared), data collection is not limited to public datasets only. Nonetheless, unique image checksums are stored in order to identify matching images. Therefore, such checksums allow users to find public images that IQMs and/or ratings derive from. The presented resource is envisioned to train automatic QC tools and to develop human expert training programs.

**Figure 1.**
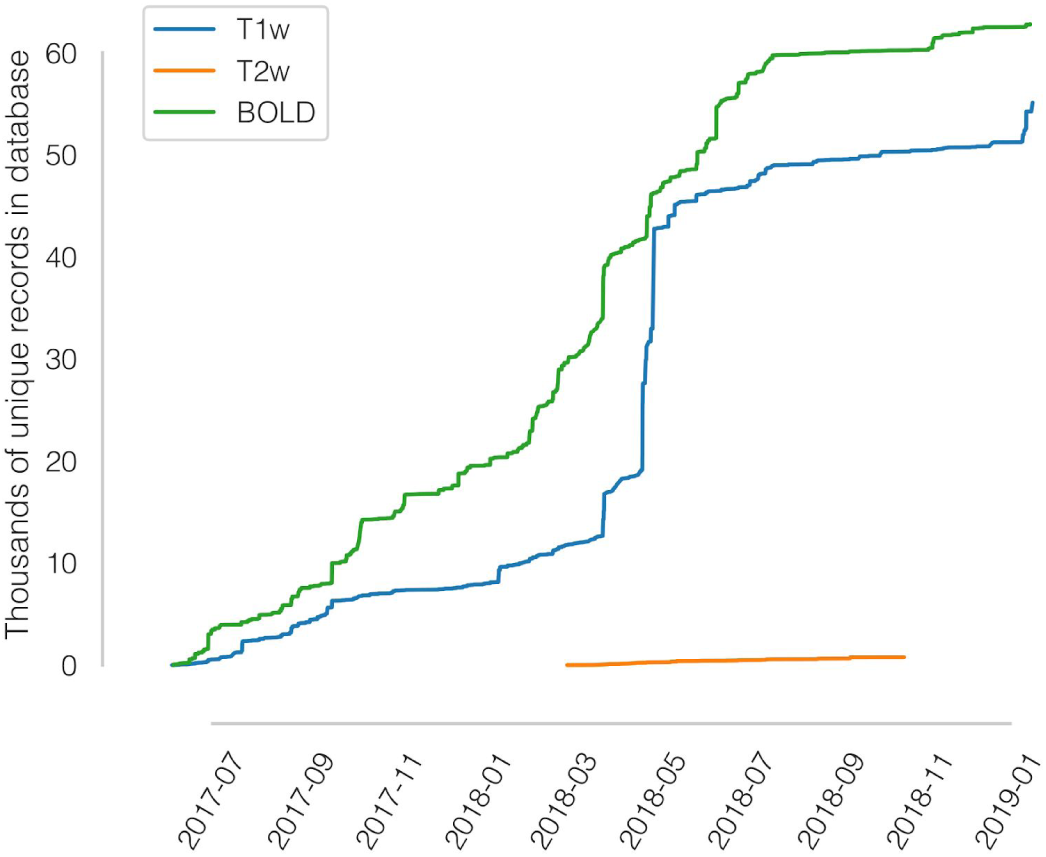
A rapidly growing MRI quality control knowledge base. The database has accumulated over 50,000 records of IQMs generated for T1-weighted (T1w) images and 60,000 records for BOLD images. Records presented are unique, i.e. after exclusion of duplicated images.

**Figure 2.**
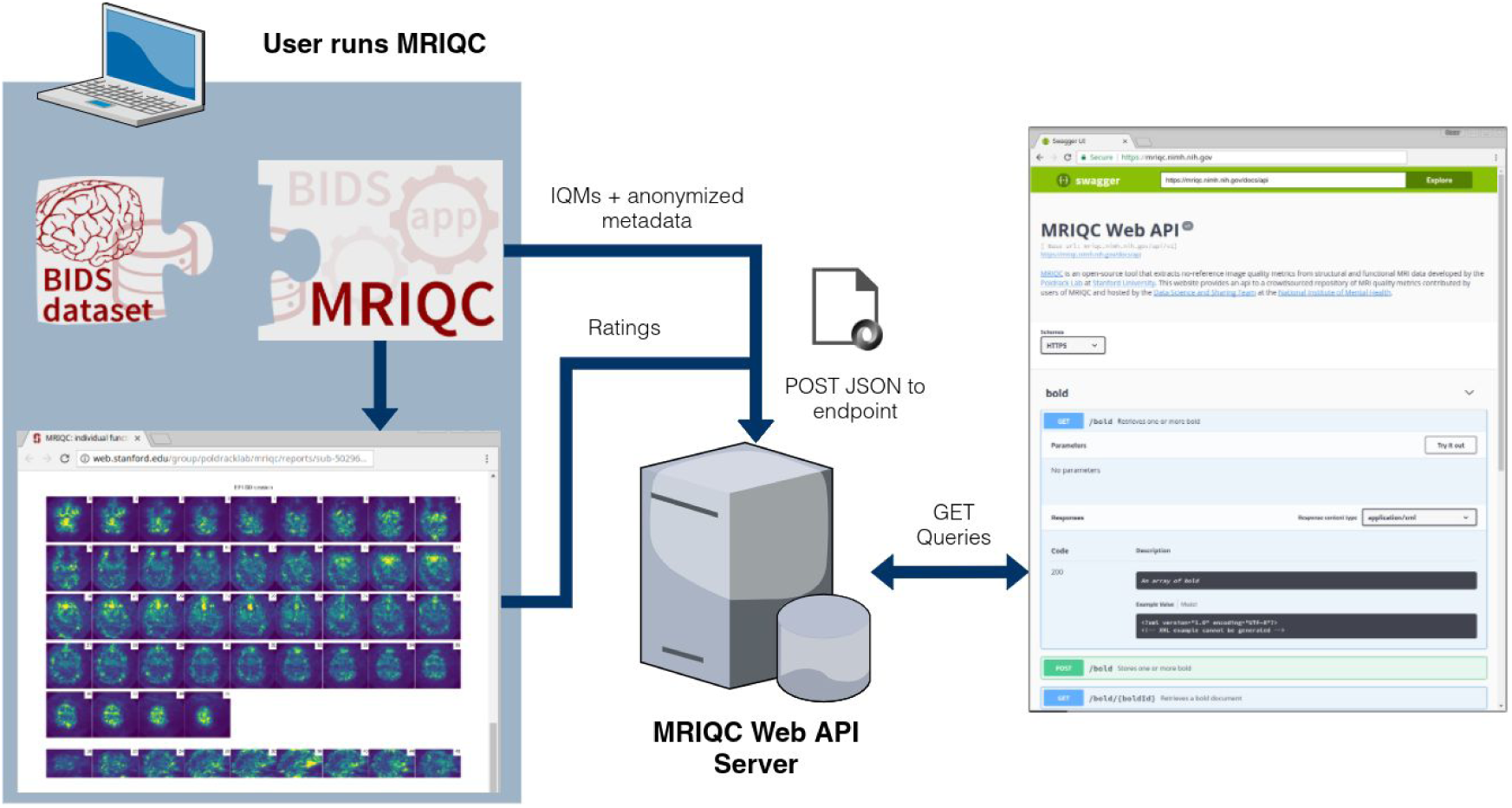
Experimental workflow to generate the database. A dataset is processed with MRIQC. Processing finishes with a POST request to the MRIQC Web API endpoint with a payload containing the image quality metrics (IQMs) and some anonymized metadata (e.g. imaging parameters, the unique identifier for the image data, etc.) in JSON format. Once stored, the endpoint can be queried to fetch the crowdsourced IQMs. Finally, a widget (Figure 3) allows the user to annotate existing records in the MRIQC Web API.

**Figure 3.**
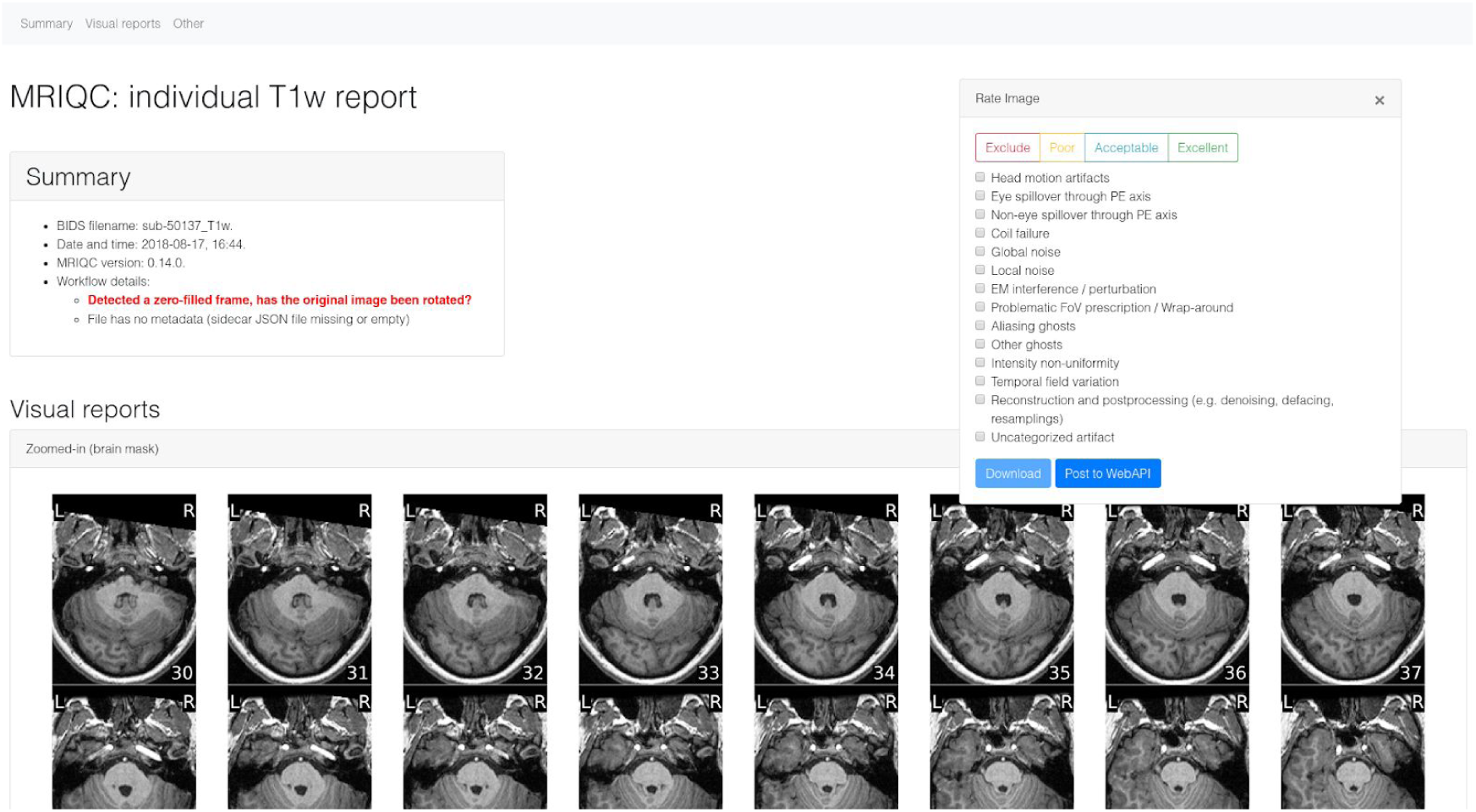
MRIQC visual reports and feedback tool. The visual reports generated with MRIQC include the “Rate Image” widget. After screening of the particular dataset, the expert can assign one quality level (among “exclude”, “poor”, “acceptable”, and “excellent”) and also select from a list of MR artifacts typically found in MRI datasets. When the annotation is finished, the user can download the ratings to their local hard disk and submit them to the Web API.

## Methods

Here we describe an open database that collects both IQM vectors extracted from functional and anatomical MRI scans, along with quality assessments done by experts based on visual inspection of images. Although it was envisioned as a lightweight web-service tailored to MRIQC, the database is able to receive new records from any other software, provided they are able to correctly query the API (application programming interface).

### Data generation and collection workflow

The overall framework involves the following workflow (summarized in Figure 2):

1. Execution of MRIQC and submission of IQMs: T1w, T2w, and BOLD images are processed with MRIQC, which computes a number of IQMs (described in section Technical Validation). The IQMs and corresponding metadata are formatted in JavaScript Object Notation (JSON), and MRIQC automatically submits them to a representational state transfer (REST) or RESTful endpoint of the Web-API. Users can opt-out if they do not wish to share their IQMs.
2. JSON records are received by the endpoint, validated, and stored in the database. Each record includes the vector of IQMs, a unique checksum calculated on the original image, and additional anonymized metadata and provenance.
3. Visualization of the individual reports: MRIQC generates dynamic HTML (hypertext markup language) reports that speed up the visual assessment of each image of the dataset. Since its version 0.12.2, MRIQC includes a widget (see Figure 2) that allows the researcher to assign a quality rating to the image being screened (see Table 3).
4. Crowdsourcing expert quality ratings: the RESTful endpoint receives the quality ratings, which are linked to the original image via their unique identifier.
5. Retrieving records: the database can be queried for records with any HTTP (HyperText Transfer Protocol) client or via the web using our interface: https://mriqc.nimh.nih.gov/. Additionally, a snapshot of the database at the time of writing has been deposited to FigShare^20^.

### Code availability

The MRIQC Web API is available under the Apache-2.0 license. The source code is accessible through GitHub (https://github.com/poldracklab/mriqcwebapi).

MRIQC is one possible client to generate IQMs and submit rating feedback. It is available under the BSD 3-clause license. The source code is publicly accessible through GitHub (https://github.com/poldracklab/mriqc).

## Data Records

A full data snapshot of the database at the time of submission is available at FigShare^20^. Alternatively, data are accessible via DataLad^21^ with the dataset URL https://github.com/oesteban/mriqc-webapi-snapshot. Table 1 describes the structure of the dataset being released.

**Table 1.** List of data tables retrieved from MRIQC-WebAPI. The following datasets are available at FigShare^20^. The <name>_curated.csv file versions correspond to the original tables after matching checksums to images in publicly available databases (and further curation as shown in https://www.kaggle.com/chrisfilo/mriqc-data-cleaning).

| Filename | Size | Description |
| --- | --- | --- |
| bold.csv | 71MB | IQMs and metadata of BOLD images (unique records) |
| bold_curated.csv | 162MB | Same as bold.csv, after curation and checksum matching |
| T1w.csv | 79MB | IQMs and metadata of T1w images (unique records) |
| T1w_curated.csv | 110MB | Same as T1w.csv, after curation and checksum matching |
| T2w.csv | 1.1MB | IQMs and metadata of T2w images (unique records) |
| T2w_curated.csv | 1.7MB | Same as T2w.csv, after curation and checksum matching |
| rating.csv | 131kB | Manually assigned quality annotations |

To obtain the latest updated records, the database can be programmatically queried online to get all the currently stored records through its RESTful API.

MRIQC reports, generated for all T1w images found in OpenfMRI are available for expert training at https://mriqc.s3.amazonaws.com/index.html#openfmri/.

## Technical Validation and Limitations

MRIQC extends the list of IQMs from the quality assessment protocol^10^ (QAP), which was constructed from a careful review of the MRI and medical imaging literature. The technical validity of measurements stored to the database is demonstrated by our previous work^9^ on the MRIQC client tool and its documentation website: https://mriqc.readthedocs.io/en/latest/measures.html. Definitions for the anatomical IQMs are given in Table 2, and for functional IQMs in Table 3. Finally, the structure of data records containing the manual QC feedback is summarized in Table 4.

**Table 2.**
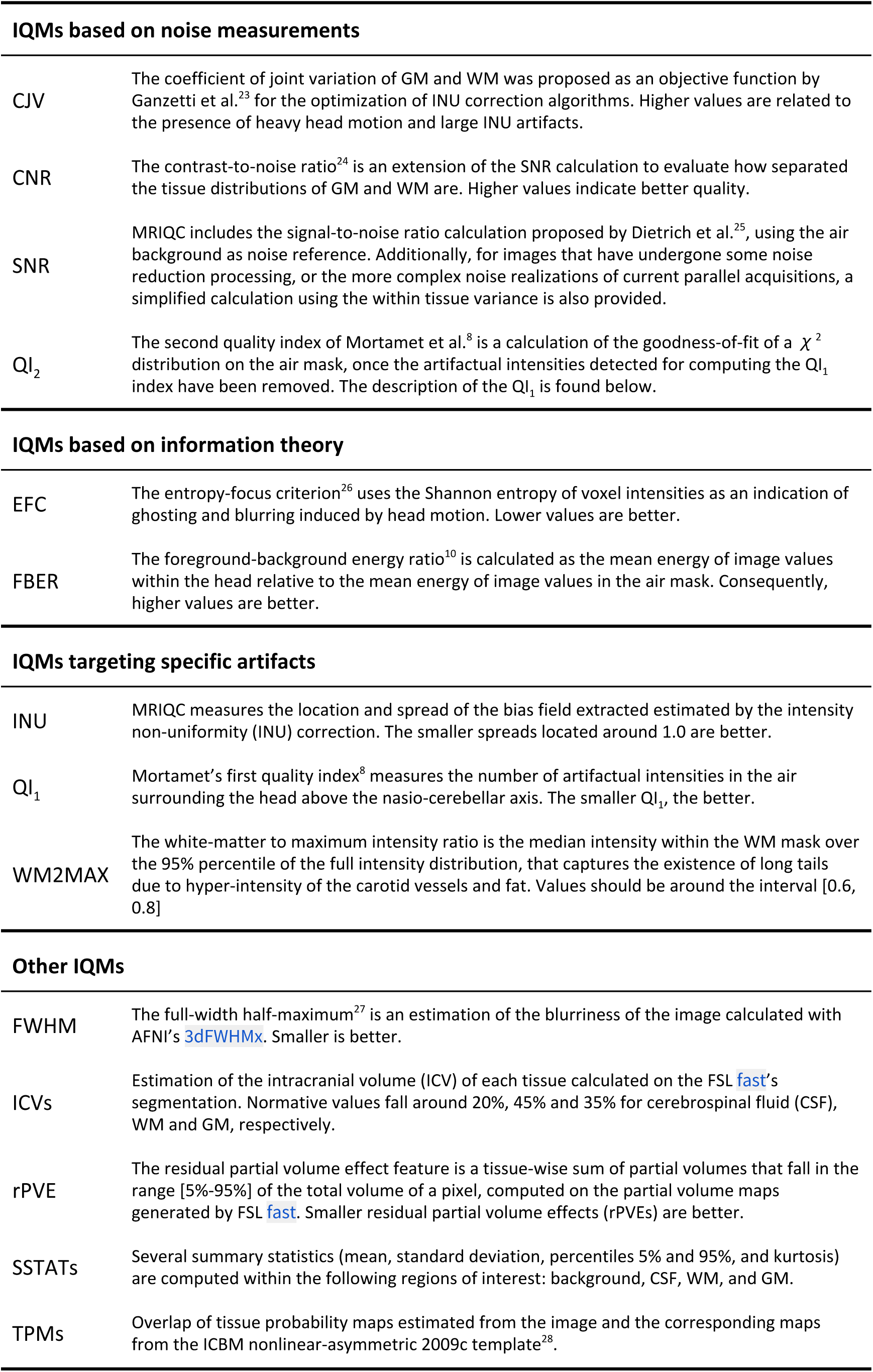
Summary table of image quality metrics for anatomical (T1w, T2w) MRI. MRIQC produces a vector of 64 image quality metrics (IQMs) per input T1w or T2w scan. (Reproduced from https://doi.org/10.1371/journal.pone.0184661.t002)

**Table 3.** Summary table of image quality metrics for functional (BOLD) MRI. MRIQC produces a vector of 64 image quality metrics (IQMs) per input BOLD scan.

| Spatial IQMs |  |
| --- | --- |
| EFC, FBER, FWHM, SNR, SSTATs (see Table 2) |  |
| IQMs measuring temporal variations |  |
| tSNR | A simplified interpretation of the original temporal SNR definition by Krüger et al.<sup>29</sup>. We report the median value of the tSNR map calculated as the average BOLD signal across time over the corresponding temporal s.d. map. |
| GCOR | Summary of time-series correlation as in<sup>30</sup> using AFNI's @compute_gcor |
| DVARS | The spatial standard deviation of the data after temporal differencing. Indexes the rate of change of BOLD signal across the entire brain at each frame of data. DVARS is calculated using *Nipype*, after head-motion correction |
| IQMs targeting specific artifacts |  |
| FD | Framewise Displacement - Proposed by Power et al.<sup>1</sup> to regress out instantaneous head-motion in fMRI studies. MRIQC reports the average FD. |
| GSR | The Ghost to Signal Ratio<sup>31</sup> estimates the mean signal in the areas of the image that are prone to N/2 ghosts on the phase encoding direction with respect to the mean signal within the brain mask<sup>10</sup>. Lower values are better. |
| DUMMY | The number of dummy scans - A number of volumes at the beginning of the fMRI time-series identified as nonsteady states. |
| IQMs from AFNI |  |
| AOR | AFNI's outlier ratio - Mean fraction of outliers per fMRI volume as given by AFNI's 3dToutcount |
| AQI | AFNI's quality index - Mean quality index as computed by AFNI's 3dTqual |

**Table 4.** Summary table of quality assessment values. Annotations received through the feedback widget are stored in a separate database collecting one rating value and an array of artifacts present in the image.

| Expert rating |  |
| --- | --- |
| Exclude | Assigned to images that show quality defects that preclude any type of processing |
| Poor | Assigned to images that, although presenting some quality problem, may tolerate some types of processing. For instance, a T1w image that may be used as the co-registration reference, but will probably generate biased cortical thickness measurements. |
| Acceptable | Assigned to images that do not show any substantial issue that may preclude processing |
| Excellent | Assigned to images without quality issues |
| Artifacts |  |
| A vector of boolean values corresponding to the following list of possible artifacts found in the image: |  |
| • Head motion artifacts<br>• Eye spillover through phase-encoding axis<br>• Non-eye spillover through phase-encoding axis<br>• Coil failure<br>• Global noise<br>• Local noise<br>• Electromagnetic interference / perturbation<br>• Problematic field-of-view prescription / Wrap-around<br>• Aliasing ghosts<br>• Other ghosts<br>• Intensity non-uniformity<br>• Temporal field variation<br>• Reconstruction and postprocessing (e.g. denoising, defacing, resamplings)<br>• Uncategorized artifact |  |

The main limitation of the database resides in that a substantial fraction of the records (e.g., around 50% for the BOLD IQMs) miss important information about imaging parameters. The original cause is that such information was not encoded with the input dataset being fed into MRIQC. However, as BIDS is permeating the current neuroimaging workflow we can expect BIDS datasets to become more complete, thereby allowing MRIQC to submit such valuable information to the Web API. Moreover, the gradual adoption of better DICOM-to-BIDS conversion tools such as HeuDiConv^22^, which automatically encodes all relevant fields in the BIDS structure, will surely help minimize this issue.

During the peer-review process of this manuscript, one reviewer identified a potential problem casting float numbers into integers on the content of the "bids_MagneticFieldStrength" field of all records. The bug was confirmed and consequently fixed on the MRIQC Web-API, and all records available on the database snapshot deposited at FigShare have been amended. When retrieving records directly from the Web-API, beware that those with creation date prior to Jan 16, 2019 require a revision of the tainted field.

## Usage Notes

Primarily, the database was envisioned to address three use-cases:

1. Sampling the distribution of IQMs and imaging parameters across datasets (including both publicly available and private), and across scanning sites.
2. Ease the image QC process, crowdsourcing its outcomes.
3. Training machines and humans.

These potential usages are revised with finer detail in the following. Note this resource is focused on quality control (QC), rather than quality assessment (QA). While QC focuses on flagging images that may endanger downstream analysis for their bad quality (i.e., identifying outliers), QA identifies issues that degrade all image's quality (i.e., improving the overall quality of images after a problem spotted in the scanning device or acquisition protocol -via QC of actual images-is fixed).

### Collecting IQMs and imaging parameters

Based on this information, researchers can explore questions such as the relationship of particular imaging parameters (e.g. MR scan vendor, or more interestingly, the multi-band acceleration factor or newest functional MRI sequences) with respect to the signal-to-noise ratio or the power of N/2 aliasing ghosts. Jupyter notebooks demonstrating examples of this use-case are available at https://www.kaggle.com/chrisfilo/mriqc/kernels.

### Crowdsourcing an optimized assessment process

To our knowledge, the community lacks a large database of multi-site MRI annotated for quality that permits the application of machine learning techniques to automate QC. As Keshavan et al. have demonstrated, minimizing the time cost and fatigue load along with the elimination of bookkeeping tasks in the quality assessment of individual MR scans enables collection and annotation of massive datasets. The graphical user interface for this use-case is presented in Figure 2.

### A database to train machines and humans

#### Training machines

As introduced before, the major bottleneck in training models that can predict a quality score for an image or identify specific artifacts, without problems to generalize across MR scanners and sites, is the small size of existing datasets with corresponding quality annotations. Additionally, these annotations, if they exist, are done with extremely varying protocols. Thus, the ability of the presented database to crowdsource quality ratings assigned by humans after visual inspection addresses both problems. The availability of multi-site, large samples with crowdsourced quality annotations that followed a homogeneous protocol (the MRIQC reports) will allow building models that overperform the random forests classifier of MRIQC^9^, in the task of predicting the quality rating a human would have assigned to an image, given a vector of IQMs (i.e., from IQMs to quality labels). Matching public image checksums, this resource will also enable to train end-to-end (from images to quality labels) deep-learning solutions

#### Training humans

Institutions can use the resource to train their experts and compare their assessments across themselves and against the existing quality annotations corresponding to publicly available datasets. Programs for training experts on quality assessment can be designed to leverage the knowledge shared via the proposed database.

## Acknowledgments

This work was supported by the Laura and John Arnold Foundation (R.A.P. and K.J.G.), the NIH (grant NBIB R01EB020740, Satrajit S. Ghosh), NIMH (R24MH114705 and R24MH117179, R.A.P; ZICMH002884, Peter Bandettini; and ZICMH002960, A.G.T.), and NINDS (U01NS103780, R.A.P.).

## Author contributions

OE - Data curation, investigation, software, validation, visualization, writing & editing.

RWB - Conceptualization, investigation, software, validation, visualization, writing & editing. DMN - Data curation, investigation, infrastructure, software, validation, writing & editing.

JCV - Investigation, infrastructure, software, validation, writing & editing.

SM - Funding acquisition, infrastructure, resources, supervision, writing & editing. AGT - Funding acquisition, infrastructure, resources, supervision, writing & editing. RAP - Funding acquisition, resources, supervision, writing & editing.

KJG - Conceptualization, investigation, software, validation, resources, supervision, writing & editing.

